# Aggression and discrimination among closely versus distantly related species of *Drosophila*

**DOI:** 10.1101/438374

**Authors:** Tarun Gupta, Sarah E. Howe, Marlo L. Zorman, Brent L. Lockwood

**Author notes:** **Author for correspondence:** Brent L. Lockwood. **Data availability:** Raw data from the scoring of aggression videos is available from the authors upon request. Representative video clips of fights are included as Supplementary Information. **Competing Interests:** The authors declare no competing interests.

## Abstract

Fighting between different species is widespread in the animal kingdom, yet this phenomenon has been relatively understudied in the field of aggression research. Particularly lacking are studies that test the effect of genetic distance, or relatedness, on aggressive behavior between species. Here we characterized male-male aggression within and between species of fruit flies across the *Drosophila* phylogeny. We show that male *Drosophila* discriminate between conspecifics and heterospecifics and show a bias for the target of aggression that depends on the genetic relatedness of opponent males. Specifically, males of closely related species treated conspecifics and heterospecifics equally, whereas males of distantly related species were overwhelmingly aggressive toward conspecifics. To our knowledge, this is the first study to quantify aggression between *Drosophila* species and to establish a behavioral bias for aggression against conspecifics versus heterospecifics. Our results suggest that future study of heterospecific aggression behavior in *Drosophila* is warranted to investigate the degree to which these trends in aggression among species extend to broader behavioral, ecological, and evolutionary contexts.

## Introduction

Heterospecific aggression—i.e., fighting between members of different species—is widespread in the animal kingdom (Peiman and Robinson, 2010). Aggressive behavior often mediates competitive interactions between species that can have important consequences for species coexistence and the structure of ecological communities (Kishi, 2015; Pfennig and Pfennig, 2012; Violle et al., 2011; Weber and Strauss, 2016). Yet, most research into aggressive behavior has focused on conspecific aggression—i.e., fighting between members of the same species (Grether et al., 2013)—with few well-characterized examples of heterospecific aggression (Peiman and Robinson, 2007), particularly in a broad phylogenetic context (Grether et al., 2017).

Fruit flies in the genus *Drosophila* present a unique opportunity to investigate aggressive behaviors, both within and between species and in a broad phylogenetic context. There are approximately 1,500 described species of *Drosophila*, many of which overlap spatially and temporally (Nielsen and Hoffmann, 1985; Prabhakaran and Sheeba, 2012; Yukilevich, 2012) and utilize similar territories and food res ources for feeding, breeding and ovipositing (da Cunha et al., 1951). Yet, while it is well established that *Drosophila* use aggression within species to establish territories and social dominance and to compete for mates and food resources (Baxter et al., 2015; Dow and von Schilcher, 1975; Hoffmann, 1987; Hoffmann and Cacoyianni, 1990; Sturtevant, 1915; White and Rundle, 2015), heterospecific aggression is largely uncharacterized, except for limited qualitative observations of heterospecific aggression among the Hawaiian *Drosophila* (Spieth, 1981).

Here we characterized male-male aggression in *Drosophila* in a multi-species context using a behavioral choice assay, in order to (1) quantify the extent to which male *Drosophila* discriminate between conspecifics and heterospecifics during aggressive social interactions and (2) test the effect of phylogenetic distance between opponent species on the distributional bias in aggressive targeting (heterospecific vs. conspecific). We report that males showed significant bias in aggression toward either conspecifics or heterospecifics in a majority of species-species interactions. Among species pairs that were more distantly related, the direction of aggression was biased toward conspecifics, whereas closely related species treated conspecifics and heterospecifics equally. To our knowledge, this is the first study to quantify aggression between *Drosophila* species and to establish a behavioral bias for aggression against conspecific vs. heterospecific opponents.

## Materials and Methods

### *Drosophila* species and husbandry

Seven species were selected from the *ananassae*, *melanogaster*, and *pseudoobscura* subgroups within the subgenus *Sophophora* (Fig. 1). Among these seven species, we assayed aggressive interactions between two species at-a-time for a total of six species pairs. Three of these species pairs are relatively closely related sibling species: i) *D. ananassae* and *D. pallidosa*, ii) *D. melanogaster* and *D. simulans*, and iii) *D. pseudoobscura* and *D. persimilis*. Whereas the other three species pairs are more distantly related: i) *D. ananassae* and *D. atripex*, ii) *D. ananassae* and *D. melanogaster*, and iii) *D. ananassae* and *D. simulans* (Fig. 1). All seven species have broad geographic distributions, and for each species pair the geographic distributions overlap (Markow and O’Grady, 2006). To the best of our knowledge, none of these species exhibit lekking behavior. Male-male aggression has previously been documented in *D. melanogaster* and *D. simulans* (Hoffmann, 1987; Kravitz and de la Paz Fernandez, 2015), but not in the other 5 species. We used one isogenic (isofemale) line from each species that was originally established from wild collections.

**Figure 1.**
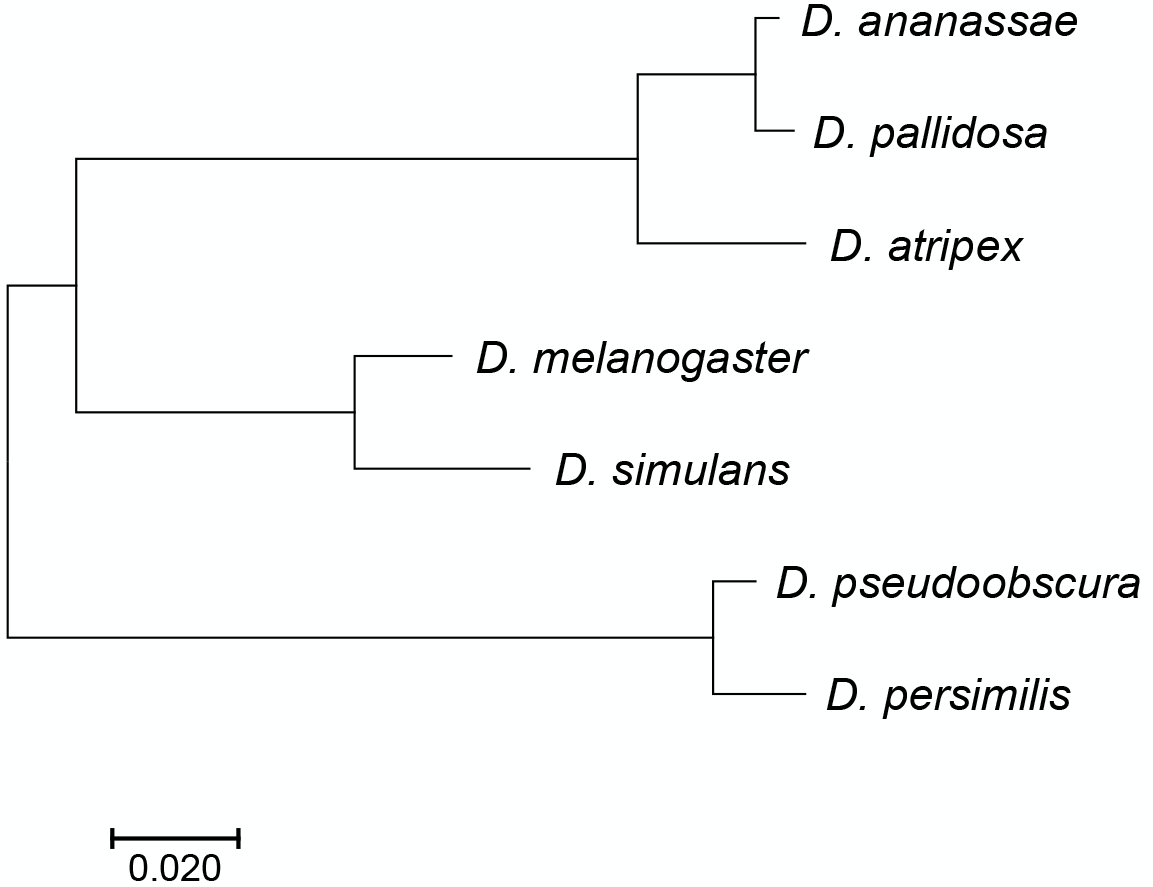
Phylogenetic relationships among focal species. Branch lengths reflect the number of substitutions per nucleotide site, as indicated by the scale bar.

*D. melanogaster* was obtained from the Bloomington Stock Center (Bloomington, IN; Canton-S; Stock No.: 64349). The *D. simulans* stock was generously provided by Brandon Cooper and Michael Turelli. The following stocks were obtained from the *Drosophila* Species Stock Center (San Diego, CA, USA): *D. ananassae* (14024-0371.15), *D. pallidosa* (14024-0433.00), *D. atripex* (14024- 0361.03), *D. pseudoobscura* (14011-0121.148), and *D. persimilis* (14011-0111.50). All flies were reared on cornmeal-based fly food containing: 1% (w/v) agarose, 8% (w/v) yeast, 4.5% molasses, and 10% (w/v) cornmeal in standard 25 mm × 95 mm polystyrene vials under humidity and temperature controlled conditions (25°C, 50% humidity and 12:12 hr light-dark cycles). *D. pseudoobscura* and *D. persimilis* were reared at 19°C, 50% humidity and 12:12 hr light-dark cycles.

### Aggression assay

To quantify agonistic social interactions in a multi-species context, we used a slightly modified version of the standard dyadic aggression assay (Mundiyanapurath et al., 2007). For each species pair, aggressive behaviors were quantified by placing two socially naïve adult males from each opponent species–a total of four males–in a standard aggression arena (Fig. 2) and measuring (1) the delay to onset of aggression (latency to aggression) and (2) the total number of aggressive lunges–a key indicator of aggression (Kravitz and de la Paz Fernandez, 2015)–by each male towards both conspecific and heterospecific opponents. We examined a total of twelve sets of interactions, as we tracked aggressive behaviors for each focal species across six species pairs. In contrast to dyadic assays typically employed in aggression studies in *Drosophila* (Certel and Kravitz, 2012; Chen et al., 2002), our multi-individual, multi-species paradigm allows examination of social behavior in a context where multiple individuals from different species compete for shared resources or territory, and it also allows us to quantify choice behavior—i.e., bias in aggression toward heterospecifics vs. conspecifics. Male pupae were isolated in 16 ×100 mm borosilicate glass tubes containing 1.5 ml of standard food medium and aged individually for 3-4 days to prevent social conditioning or formation of social dominance hierarchies prior to testing. Three-day old adult males were extracted under CO_2_ anesthesia and marked on the thorax with a dab of white or blue acrylic paint (assigned randomly) for species identification during assay setup and scoring. After painting, males were transferred to new isolation tubes containing 1.5 ml agarose-based nutritionally deficient media (without cornmeal, yeast or sugar) and allowed recovery from handling and anesthesia. The following day, two 4-5 day old, socially naïve adult males from each opponent group – a total of 4 males – were gently aspirated into one of the wells of a 12-well polystyrene plate (Thermo Fisher Scientific #130185) with a small cup in the middle containing food – representing focal point of contest (Fig. 2). All four males were introduced to the chamber at the same time to prevent a potential resident-intruder confound. All behavioral assays were set up and recorded within 0-2 Zeitgeber hours, i.e., the first two hours of the lights ON time in a 12:12 light-dark cycle.

**Figure 2.**
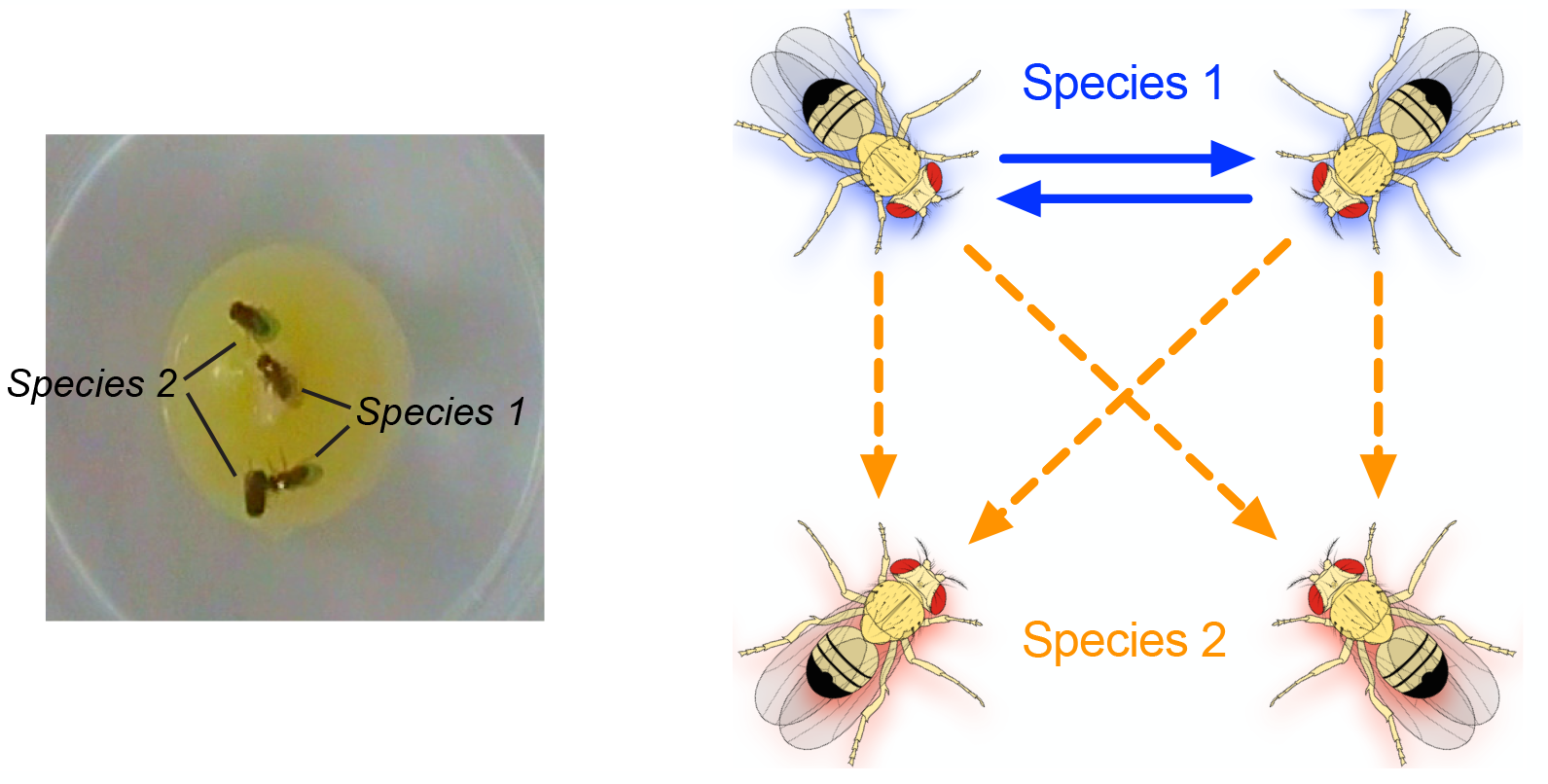
Multi-species aggression assay. Left panel is an image of the aggression arena showing separate species labeled on the thoraxes by white paint (visible in the image, species 1) or blue paint (not visible in the image, species 2). Right panel illustrates the null expectation of a 1:2 ratio of the number of conspecific lunges to heterospecific lunges.

### Aggression scoring

The number of lunges against conspecific and heterospecific males was counted for a period of 30- minutes after the first lunge, for consistency with the scoring duration of aggression assays reported elsewhere (Certel and Kravitz, 2012; Chen et al., 2002). The amount of time between the introduction of males to the aggression chamber and the first aggressive lunge was used as the measurement of delay to the onset of aggression, or latency to lunge. The latency to lunge was scored separately for the two directions of lunges, conspecific or heterospecific. Scoring was terminated after 1 hour if no aggressive encounter was recorded during that period. Aggressive behaviors were scored manually by two independent scorers using iMovie ‘09, version 8.0.6 (Apple Inc., Cupertino, CA, USA). The number of aggression trials is indicated in Figures 3 and 4.

**Figure 3.**
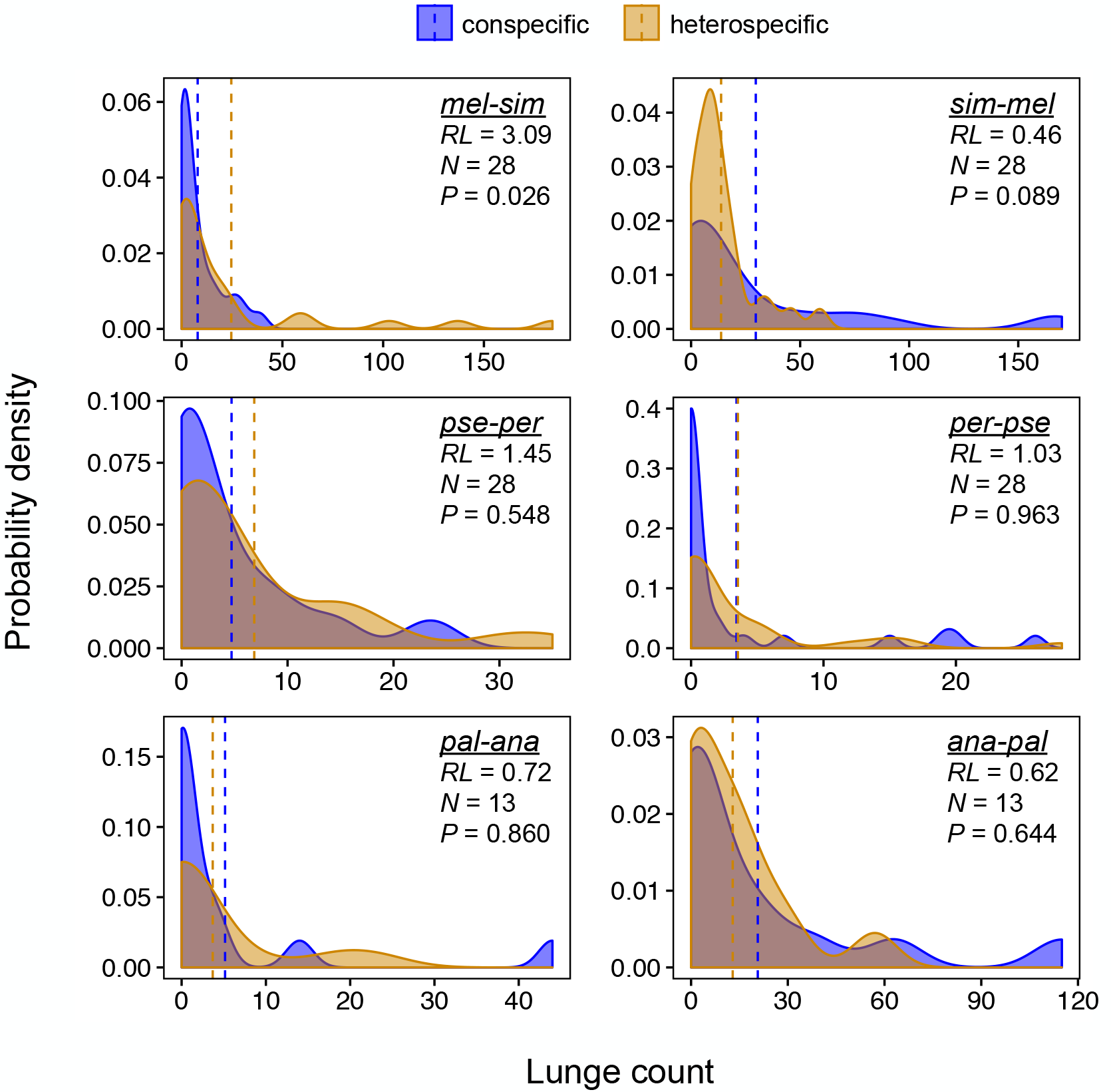
Closely related species pairs were equally aggressive toward heterospecifics and conspecifics. Shown are probability density plots of conspecific (blue) and heterospecific (orange) lunge counts of closely related species pairs. Focal species is the aggressor that lunges on itself or the non-focal species (focal-nonfocal), with the same species name abbreviations as in Table 1. Vertical dashed lines indicate mean lunge counts. Ratio of mean lunge counts (*RL*; heterospecific:conspecific), number of aggression trials (*N*), and significance of the difference between the distributions of lunge counts based on the z-test (*P*) are shown in the upper right of each panel.

**Figure 4.**
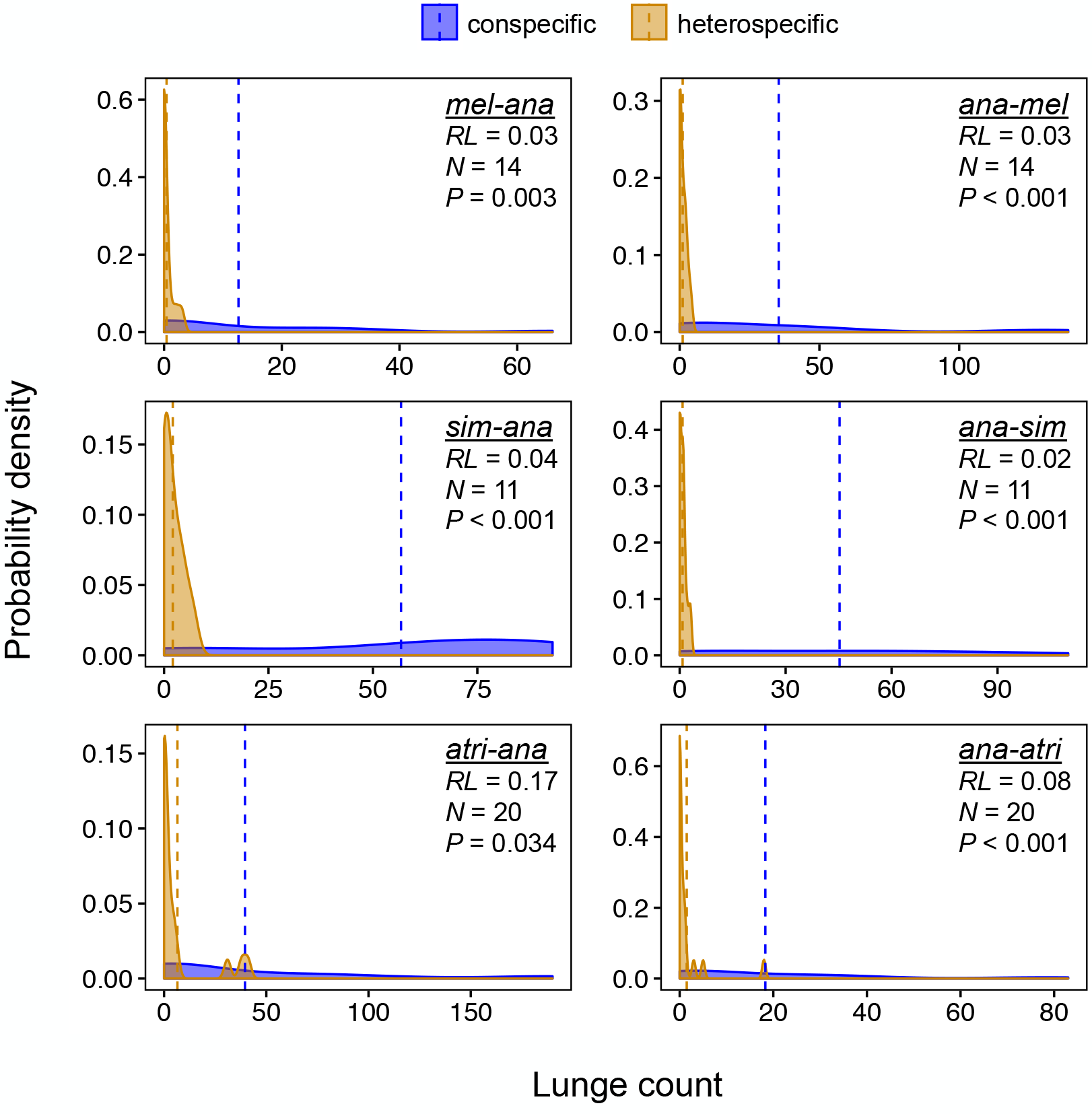
Distantly related species pairs preferentially targeted aggression at conspecifics. Shown are probability density plots of conspecific (blue) and heterospecific (orange) lunge counts of distantly related species pairs. Focal species is the aggressor that lunges on itself or the non-focal species (focal-nonfocal), with the same species name abbreviations as in Table 1. Vertical dashed lines indicate mean lunge counts. Ratio of mean lunge counts (*RL*; heterospecific:conspecific), number of aggression trials (*N*), and significance of the difference between the distributions of lunge counts based on the z-test (*P*) are shown in the upper right of each panel.

### Phylogenetic distance

We inferred evolutionary relationships (Fig. 1) and calculated pairwise genetic distances (Fig. 6) among species using the Maximum Likelihood method based on the Tamura-Nei model (Tamura and Nei, 1993) using MEGA7 (Kumar et al., 2016). We compared 3,196 nucleotide positions among 1 mitochondrial gene (*CoI*) and 2 nuclear genes (*Gpdh* and *kl2*), for which sequence data were readily available on NCBI for all species included in the study. Although full genome sequences are available for *D. melanogaster*, *D. simulans*, *D. pseudoobscura*, *D. persimilis*, and *D. ananassae*, there is relatively little nucleotide sequence data available for *D. pallidosa* and *D. atripex*.

Nonetheless, the evolutionary relationships that we report herein are consistent with previous studies (Clark et al., 2007; Matsuda et al., 2009). We downloaded sequences from NCBI of *D. pallidosa* (*CoI*, Accession #: FJ795561; *Gpdh*, FJ795596; and *kl2*, FJ795633) and *D. atripex* (*CoI*, FJ795575; *Gpdh*, FJ795601; and *kl2*, FJ795643) and used these to BLAST the published genome sequences of the other five species included in the study. The phylogenetic tree with the highest log likelihood (- 6435.05) is shown in Figure 1. Initial trees for the heuristic search were obtained by Neighbor-Join and BioNJ to a matrix of pairwise distances estimated with the Maximum Composite Likelihood approach. A discrete Gamma distribution was used to model evolutionary rate differences among sites. All positions containing gaps were removed from the analysis. Branch lengths correspond to the number of substitutions per site.

### Body-size estimation

In many species, aggressiveness correlates with body size both within and between species and smaller males are less likely to initiate and hold aggressive encounters (Hoffmann, 1987). Body length (mm) from anterior antennae to posterior abdomen of males from all opponent group used in aggression assays was measured as a proxy for body size using ImageJ (Schneider et al., 2012).

### Statistical analyses

We compared the number of lunges directed toward conspecifics vs. heterospecifics with a negative binomial generalized linear model, as implemented in the MASS package in R version 3.3.2 (R Core Team 2016). We normalized the number of heterospecific lunges by dividing by two (rounded to the nearest whole number) because there were twice as many heterospecific opponent males as conspecific opponent males in the aggression arena (Fig. 2). Goodness-of-fit was assessed by a chi-square test of the residual deviance of the negative binomial model. To examine the direction and effect size of aggression bias, we calculated the ratio of mean lunge counts (*RL*), which is the ratio of the mean number of heterospecific lunges to the mean number of conspecific lunges. *RL* was obtained by exponentiating the regression coefficient (β) of the negative binomial model, as this coefficient equals the log-ratio of mean lunge counts. The negative binomial model was also used to estimate the 95% confidence intervals of *RL*, which are reported in Figure 5. Significant differences between the distributions of heterospecific and conspecific lunge counts were determined by a z-test of the estimated regression coefficient and standard error from the negative binomial model. We ran separate regressions for each focal species in each species pair for a total of 12 regressions (6 species pairs x 2 species in each pair). *P*-values were corrected for multiple tests via the Benjamini-Hochberg method (Benjamini and Hochberg, 1995).

**Figure 5.**
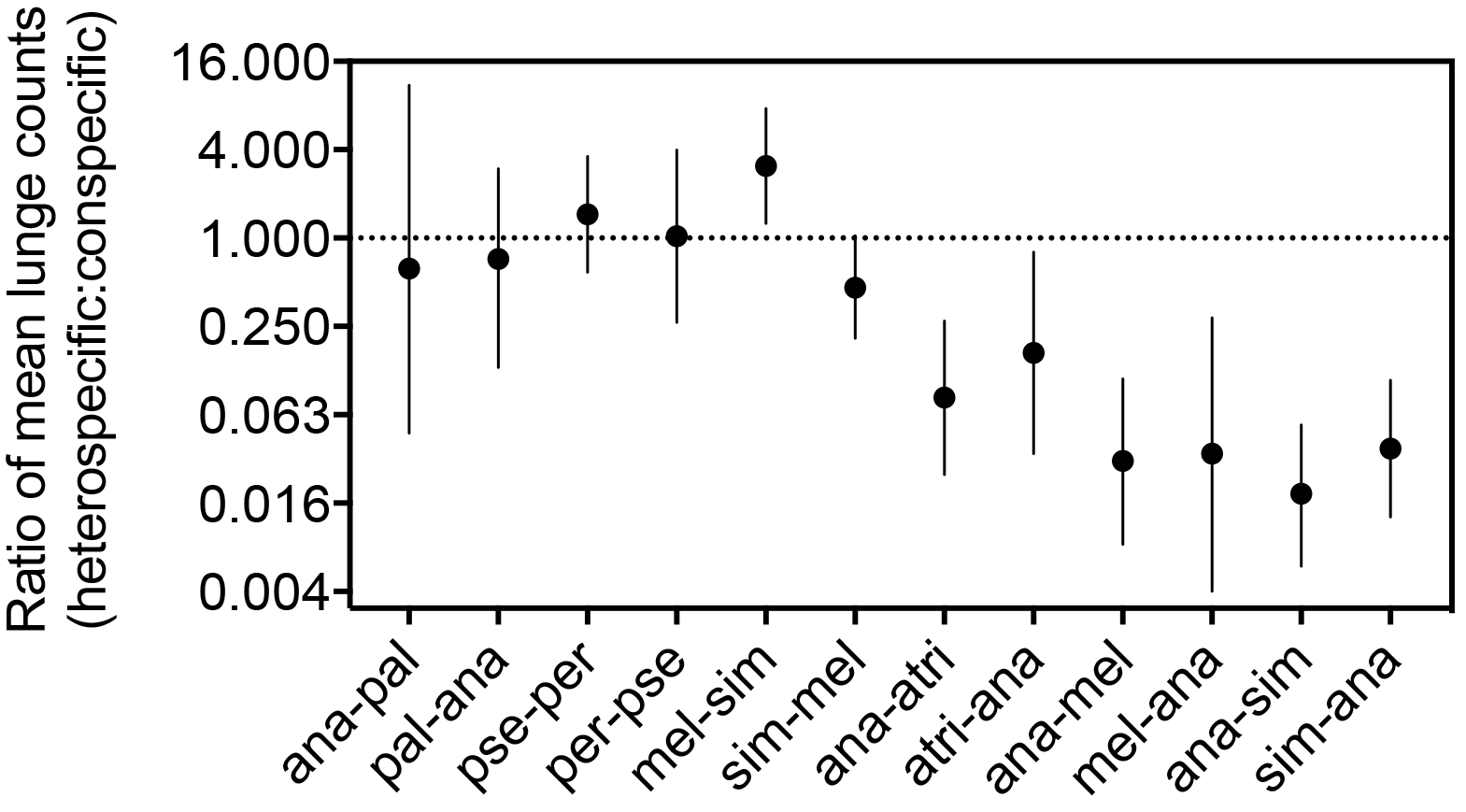
Ratio of mean lunge counts for each species pair interaction. The dotted line at 1 indicates the null expectation that heterospecifics and conspecifics were treated equally. Values greater than 1 indicate greater aggression toward heterospecifics, and values less than 1 indicate greater aggression toward conspecifics. The data are presented on a log scale. Focal species is the aggressor that lunges on itself or the non-focal species (focal-nonfocal), with the same species name abbreviations as in Table 1. Species pairs are ordered from most closely related to most distantly related (left to right). Error bars represent 95% confidence intervals.

We assessed the relationship between aggression bias (*RL*) and genetic distance between opponent species via permutation analyses. Given the design of our aggression assay and the phylogenetic relatedness of the focal species, aggression biases among species pairs may not be independent. To account for this lack of independence among data points in our analyses, we performed 10,000 permutations of the genetic distance vs. *RL* relationships—i.e., the genetic distances and *RL* values for each species pair were randomly shuffled and resampled—and significance was assessed based on the probability distribution of the Spearman rank coefficient.

We examined the relationship between the latency to lunge and the direction of the first lunge (conspecific vs. heterospecific) via a 2-way ANOVA, with species pair and direction of lunge as fixed effects. We examined the relationships between (1) body length and number of lunges from focal males and (2) body length difference between opponent species and total number of heterospecific lunges via the Spearman correlation. All analyses were conducted in R version 3.3.2 (R Core Team 2016) or GraphPad Prism version 7 (GraphPad Software, Inc., La Jolla, CA, USA).

## Results

We observed a significant distributional bias in the targets of aggression—i.e., lunges directed toward either conspecific or heterospecific opponent males—in seven out of twelve species-pair interactions (Table 1). The behavior of closely related species pairs contrasted with that of more distantly related species pairs. Among closely related species pairs, heterospecifics and conspecifics were treated more or less equally (i.e., there was not a strong bias in the direction of aggression), as can be seen in the largely overlapping distributions of heterospecific and conspecific lunge counts (Fig. 3). In addition, the ratios of mean lunge counts (*RL*; heterospecific:conspecific) in closely related species pairs hovered around values of one (Fig. 5; *RL* ≈ 1), indicating that heterospecifics and conspecifics were equally likely to be targeted by aggression. The only closely related species pair interaction that showed a significant aggression bias was *D. melanogaster* paired with *D. simulans* (Table 1), where *D. melanogaster* males were 3 times more likely to target heterospecifics than conspecifics (Figs. 3 and 5; *RL* = 3.09). In contrast, among more distantly related species pairs, the distributions of heterospecific and conspecific lunges did not overlap (Fig. 4), and the ratios of mean lunge counts were all less than one (Fig. 5; *RL* < 1), indicating strong conspecific aggression biases.

**Table 1.** Statistics of the negative binomial model fits. Bold italic indicates significant differences between the distributions of heterospecific and conspecific lunge counts after false discovery rate correction.

| Focal-nonfocal | Residual deviance | Residual d.f. | Goodness of fit ($\chi^2$ P-value) | $\beta \pm$ std. error | z | FDR-corrected P-value |
| --- | --- | --- | --- | --- | --- | --- |
| mel-sim | 63.20 | 54 | 0.1833 | **1.13 ± 0.45** | **2.49** | **0.0255** |
| sim-mel | 65.46 | 54 | 0.1364 | -0.77 ± 0.41 | -1.89 | 0.0890 |
| pse-per | 58.33 | 54 | 0.3193 | 0.37 ± 0.46 | 0.82 | 0.5484 |
| per-pse | 43.17 | 54 | 0.8545 | 0.03 ± 0.67 | 0.05 | 0.9633 |
| pal-ana | 16.40 | 24 | 0.8732 | -0.33 ± 1.24 | -0.27 | 0.8602 |
| ana-pal | 27.40 | 24 | 0.2860 | -0.47 ± 0.76 | -0.62 | 0.6443 |
| mel-ana | 18.76 | 26 | 0.8466 | **-3.38 ± 1.04** | **-3.26** | **0.0027** |
| ana-mel | 28.43 | 26 | 0.3378 | **-3.50 ± 0.65** | **-5.39** | **2.86E-07** |
| sim-ana | 25.20 | 20 | 0.1940 | **-3.30 ± 0.54** | **-6.12** | **5.53E-09** |
| ana-sim | 22.82 | 20 | 0.2977 | **-4.01 ± 0.56** | **-7.18** | **8.64E-12** |
| atri-ana | 37.25 | 38 | 0.5041 | **-1.80 ± 0.77** | **-2.33** | **0.0336** |
| ana-atri | 38.66 | 38 | 0.4396 | **-2.50 ± 0.60** | **-4.15** | **0.0001** |

Focal species is the aggressor that lunges on itself or the non-focal species (focal-nonfocal), with the following species name abbreviations: mel = *D. melanogaster*, sim = *D. simulans*, pse = *D. pseudoobscura*, per = *D. persimilis*, pal = *D. pallidosa*, ana = *D. ananassae*, atri = *D. atripex*.

Further supporting the contrast between the aggressive behaviors of closely versus distantly related species pairs, there was a significant negative relationship between the genetic distance between competing species and the ratio of mean lunge counts (Fig. 6; Spearman *ρ* = − 0.82, *N* = 10,000 permutations, *P* = 0.002). In other words, more distantly related species pairs were most aggressive to conspecifics, whereas closely related species pairs treated conspecifics and heterospecifics with equal levels of aggression. In fact, males in the distantly related *D. simulans* – *D. ananassae* species pair displayed a high degree of tolerance for heterospecific opponents sharing the food cup but escalated quickly to high-intensity lunging when confronted by conspecific opponents (Video S1). Conversely, the intensity of aggression directed towards heterospecifics was greatest in closely related species pairs, such as *D. simulans* – *D. melanogaster* (Video S2).

**Figure 6.**
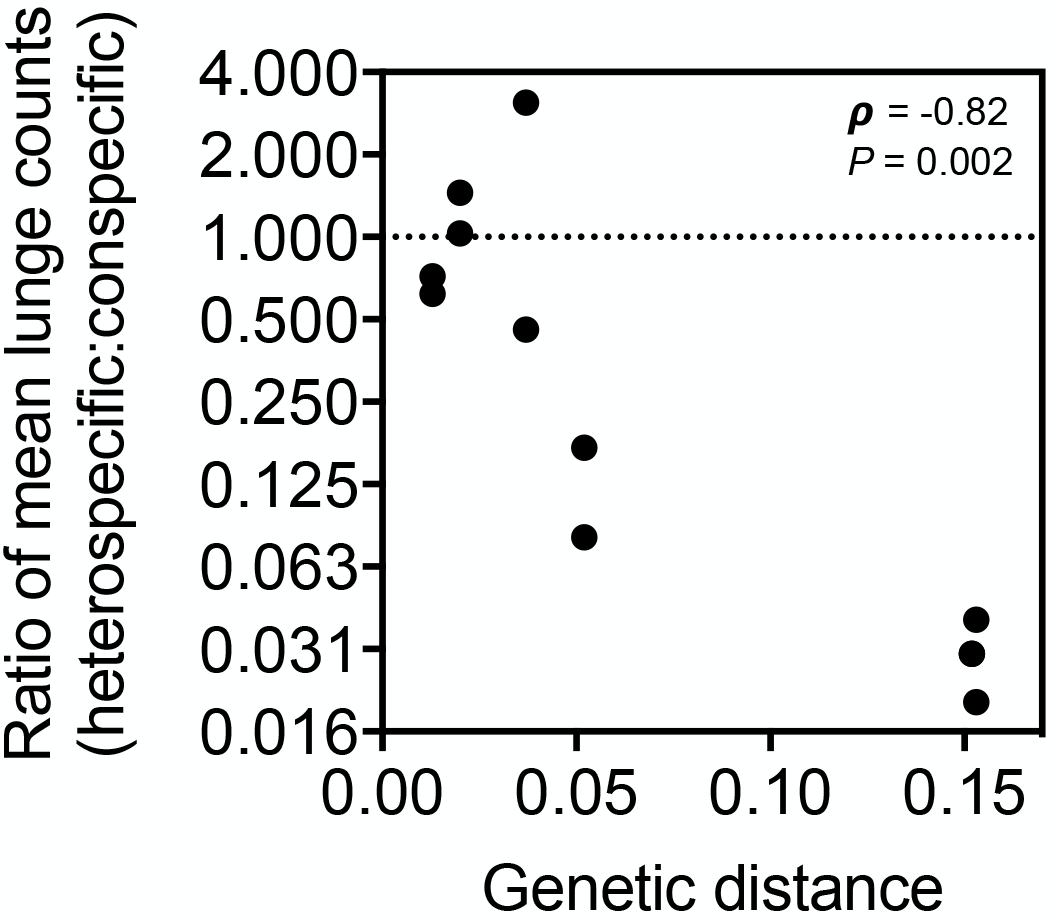
Greater genetic distance between species pairs led to significant conspecific aggression biases (Spearman *ρ* = −0.82, *N* = 10,000 permutations, *P* = 0.002). Plotted are the pairwise genetic distances versus the mean ratio of lunge counts for each species pair. The dotted line at 1 indicates the null expectation that heterospecifics and conspecifics were treated equally. Values greater than 1 indicate greater aggression toward heterospecifics, and values less than 1 indicate greater aggression toward conspecifics.

There was no significant difference in the latency to initiate aggression towards conspecifics or heterospecifics (Fig. S1; 2-way ANOVA, direction of lunge effect, *F_1,177_* = 0.00057, *P* = 0.98, direction of lunge x species pair interaction, *F_5,177_* = 1.515, *P* = 0.19). That is, males from either opponent group were equally likely to be targeted at the initial onset of aggression when low-intensity encounters first escalate to high-intensity lunging. These results suggest an opportunistic, non-selective tendency towards initiating an aggression sequence followed by a species-specific strategy for selectively targeting subsequent aggressive behaviors.

Body size differences among opponents have been shown previously to influence male fly aggressiveness (Hoffmann, 1987), but we found no significant association between average body size and number of aggressive lunges by a given species (Fig. S2B; Spearman *ρ* = 0.015, *N* = 155, *P* = 0.85). Furthermore, the relative body-size difference between opponent species in a given fight showed no significant relationship to the number of heterospecific lunges (Fig. S2C; Spearman *ρ* = −0.14, *N* = 75, *P* = 0.24).

## Discussion

To our knowledge, this is the first study to demonstrate discriminatory aggression between species of *Drosophila*—i.e., the differential aggressive response of males toward conspecifics versus heterospecifics in multi-species social interactions—albeit aggression biases were mostly observed between distantly related species and not closely related species. While males of many species of *Drosophila* are known to be territorial (Baxter et al., 2015; Chen et al., 2002; Hoffmann, 1987), particularly in the lekking species that are endemic to Hawaii (Spieth, 1974), previous work has only provided limited accounts of heterospecific interactions (Spieth, 1981), and heterospecific aggression has never been explicitly quantified.

We interpret the differential aggressive responses among closely vs. distantly related species pairs as innate responses that are mediated by species recognition cues. Because all interacting individuals in this study were extracted as pupae and socially isolated as adults with no direct contact with other males from either species, the biases in aggressive targeting are not likely to be learned behaviors. That is, unless learning occurred during the larval stages. Previously, it has been shown that *D. melanogaster* larvae learn and retain chemosensory cues from early larval stages (Durisko and Dukas, 2013). If larval learning persists through metamorphosis, then adult aggression could be influenced by social experiences or chemosensory preferences that were established during early larval stages. Therefore, we cannot rule out the possibility that closely related species pairs treated heterospecifics similarly to conspecifics simply because the chemosensory environments they were reared in were similar. If this were the case, larvae reared in isolation could potentially produce adults that behave differently than those reported on herein.

There are several potential molecular mechanisms that underlie these behavioral responses to male-male encounters of different species. Recently, it has been reported that epigenetic mechanisms, such as DNA methylation, serve as an interface between the genome and the environment and can facilitate species-specific behavioral plasticity in the context of courtship by modulating aminergic function (Gupta et al., 2017). Thus, as a proximate mechanistic cause for the bias in aggressive targeting reported in the present study, octopaminergic systems may play a critical role in relaying species-specific chemosensory information (Andrews et al., 2014), and facilitate species recognition and/or discrimination in the context of mixed-species aggressive interactions.

Based solely on the data presented herein, we cannot evaluate the ultimate evolutionary causes of male aggression biases among *Drosophila* spp. Nonetheless, it is important to consider the potential ecological and evolutionary mechanisms that influence these patterns in order to provide a framework for future work. A major outstanding question is whether these behavioral biases for aggression are due to ancestral states, where males treat closely related heterospecifics like conspecifics due to mistaken identity (i.e., falsely identifying a heterospecific opponent as a conspecific), or if aggression bias is influenced by current and ongoing interference competition. Previous studies in other animal species lend support to the mistaken identity hypothesis. In a meta-analysis of birds and fish, Peiman and Robinson (Peiman and Robinson, 2010) found that, among species that do not share resources, heterospecific aggression is greatest among closely related species. Similarly, in a separate meta-analysis of wood warbler birds, Losin et al. (Losin et al., 2016) found that, even among sympatric species, patterns of heterospecific aggression can largely be explained by shared ancestry. Thus, in many cases, heterospecific aggression may be an evolutionary artifact that originates from natural selection for conspecific aggression, which erodes over time following species divergence. In fact, it may be difficult to parse this non-adaptive cause of heterospecific aggression from the effects of interference competition between species. To overcome this challenge and potentially account for these confounding effects, Peiman and Robinson (2010) suggest comparing levels of heterospecific aggression among allopatric species vs. aggression among sympatric species. In the allopatric case, heterospecific aggression should be non-adaptive because species do not directly compete for resources. In comparison to allopatry, higher or lower levels of heterospecific aggression in sympatry could be attributed to the evolution of aggressive behaviors in response to interference competition among species.

A further consideration in analyses of heterospecific aggression, and an important caveat to the interpretation of the results we report herein, is the degree to which species pairs directly interact in nature. All of the species pairs included in this study have large biogeographic ranges that overlap to varying degrees (Markow and O’Grady, 2006), and each species pair has similar food preferences (da Cunha et al., 1951; Nielsen and Hoffmann, 1985; Prabhakaran and Sheeba, 2012). Thus, it is possible that these species pairs compete in nature, and that direct interference competition between species influences the evolution of heterospecific aggression. However, to our knowledge direct interference competition among the six species pairs included in this study has never been documented in nature. Therefore, future work is required to address whether or not these species pairs directly compete for resources in nature.

We would also like to note that direct competition for mates (i.e., reproductive interference competition) among heterospecifics is another type of interference competition that could influence heterospecific aggression. Indeed, the intensity of reproductive competition among species has been shown to be a key factor that influences heterospecific aggression in other species (Drury et al., 2015). We were not able to assess this effect in the present study because, while hybridization is known to occur in the laboratory among the closely related sibling species pairs (Yukilevich, 2012), there is lack of consensus as to whether or not hybridization occurs in nature for these species pairs (Bock, 1984; Matsuda et al., 2009; Noor, 1995; Yukilevich, 2012). Future work should focus on comparisons of taxa among sibling species of *Drosophila* that are known to hybridize in nature, such as species in the *subquinaria* complex (Dyer et al., 2014) or among the *yakuba*, *teissieri*, and *santomea* sister species (Comeault et al., 2016; Cooper et al., 2018), to examine the role of reproductive interference competition in influencing the evolution of heterospecific aggression.

## Supplemental Information

**Supplemental Figure S1.**
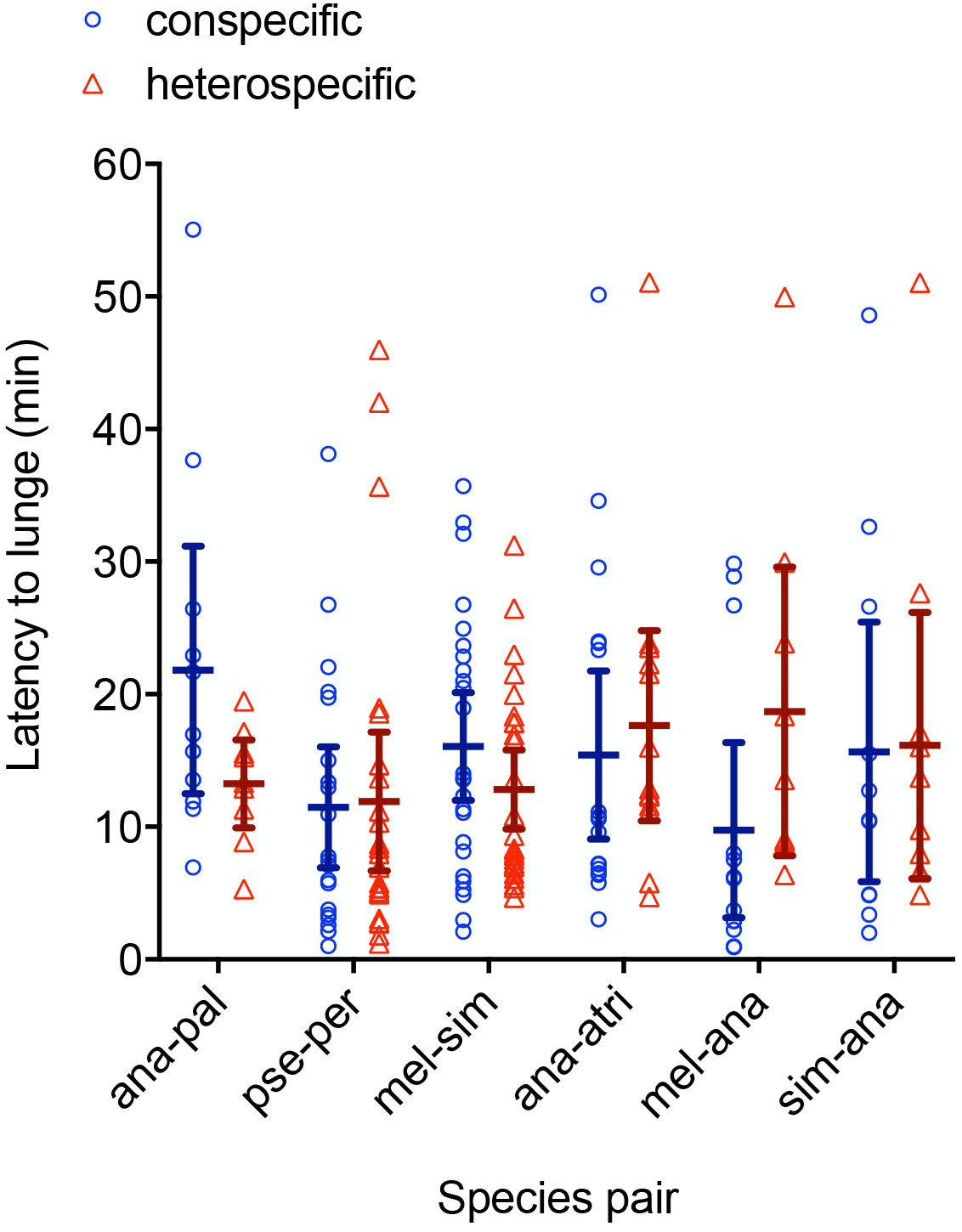
Latency to lunge is the delay to onset of aggression (minutes) following the beginning of a fight. There was no significant difference between the latencies to lunge at conspecifics vs. heterospecifics (2-way ANOVA, direction of lunge effect, *F_1,177_* = 0.00057, *P* = 0.98, direction of lunge x species pair interaction, *F_5,177_* = 1.515, *P* = 0.19), and all species pairs initiated aggression at approximately the same time (2-way ANOVA, species pair effect, *F_5,177_* = 1.088, *P* = 0.3686).

**Supplemental Figure S2.**
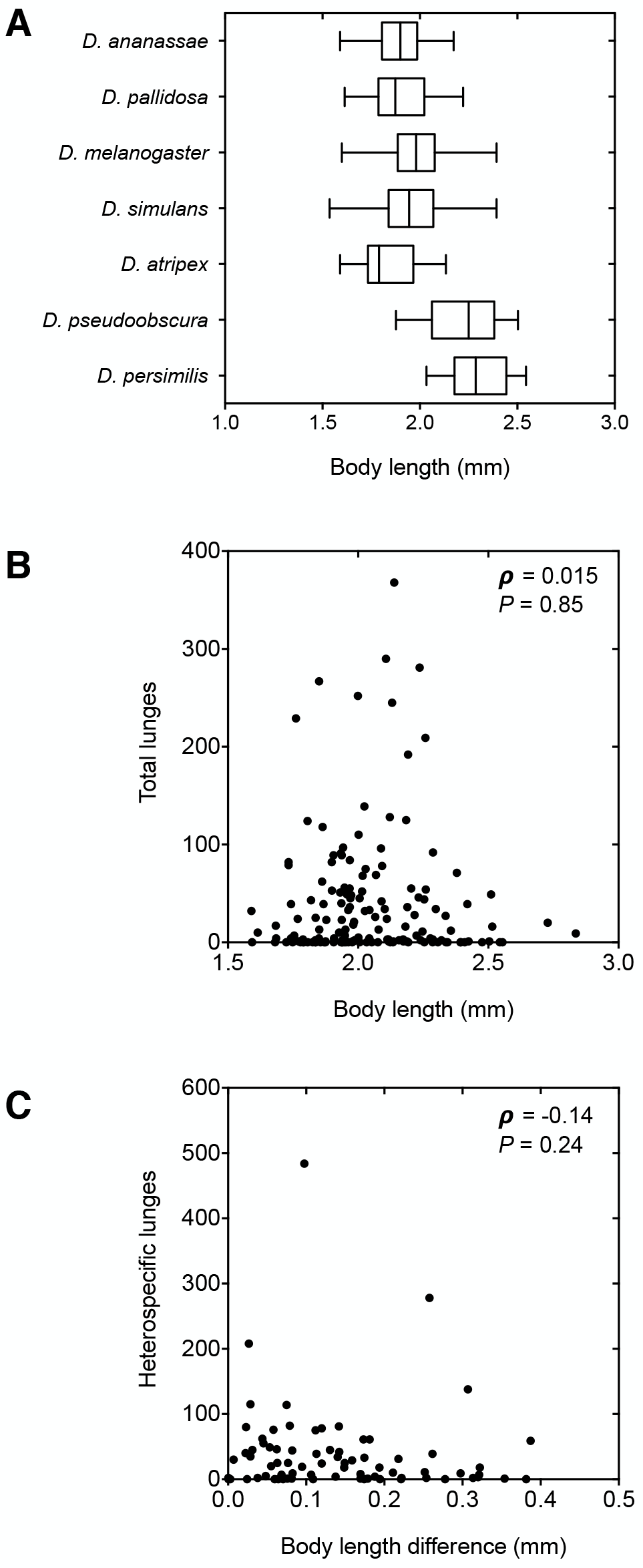
Body length and body length difference had no effect on levels of aggression or aggression bias. (*A*) Body length did not differ between opponent species (Sidak’s multiple comparisons test, d.f. = 304; *D. melanogaster* vs. *D. simulans*, *t* = 1.98, *P* = 0.26; *D. pseudoobscura* vs. *D. persimilis, t* = 1.663, *P* = 0.46; *D. pallidosa* vs. *D. ananassae, t* = 0.2431, *P* = 0.99; *D. simulans* vs. *D. ananassae, t* = 1.566, *P* = 0.53; *D. ananassae* vs. *D. atripex, t* = 1.585, *P* = 0.52), except that *D. melanogaster* was slightly larger than *D. ananassae* (mean difference in body length: 0.09 cm; Sidak’s multiple comparisons test, *t* = 3.572, *P* = 0.0025). However, size did not have an effect on aggression, as (*B*) there was no significant relationship between body length and total number of lunges from focal males in a given fight (Spearman *ρ* = −0.015, *P* = 0.85) and (*C*) there was no significant relationship between the difference in body lengths between males of opponent species and the number of heterospecific lunges in a given fight (Spearman *ρ* = −0.14, *P* = 0.24).

**Supplemental videos (Video S1 and S2) have been uploaded as separate files.**

**Supplemental Video S1.** Males of distantly related *Drosophila simulans* (white dot on thorax) and *Drosophila ananassae* (blue dot on thorax, not visible in this video) in the multi-species aggression arena. Note how nearly all aggressive encounters occur between members of the same species.

**Supplemental Video S2.** Males of closely related *Drosophila melanogaster* (white dot on thorax) and *Drosophila simulans* (blue dot on thorax, not visible in this video) in the multi-species aggression arena. Note how aggressive encounters between members of the opposite species occur at a similar frequency to aggressive encounters between members of the same species.

**Author contributions**
TG and BLL conceived and planned the study. TG, SEH, and MLZ conducted the experiments. TG and BLL analyzed the data. TG and BLL wrote the manuscript.

## Acknowledgements

We thank Brandon Cooper and Michael Turelli for generously providing the *D. simulans* stock. We thank Sara Helms Cahan, Nicholas Gotelli, Melissa Pespeni, Emily Mikucki, Dean Castillo, Leonie Moyle, and three anonymous reviewers for their helpful discussions and comments on this manuscript. This work was supported by the University of Vermont.

